## Supplemental Inforamation for "Action mechanism of a novel agrichemical quinofumelin against *Fusarium graminearum*"

**Table S1. GO analysis of down- and up-regulated DEGs.**

| Description | Up | Down |
| --- | --- | --- |
| Vitamin binding | 0 | 5 |
| Transition metal ion binding | 3 | 10 |
| Tetrapyrrole binding | 3 | 3 |
| RNA biosynthetic process | 2 | 4 |
| Phosphopantetheine binding | 0 | 4 |
| Peroxidase activity | 1 | 1 |
| Oxidoreductase activity | 1 | 11 |
| Nucleic acid-templated transcription | 4 | 8 |
| NADP binding | 0 | 3 |
| Monooxygenase activity | 1 | 6 |
| Monocarboxylic acid metabolic process | 1 | 2 |
| Modified amino acid binding | 0 | 4 |
| Homeostatic process | 1 | 1 |
| Heme binding | 3 | 3 |
| Chemical homeostasis | 1 | 1 |
| Cellular amino acid catabolic process | 0 | 2 |
| Amide binding | 0 | 4 |

**Table S2. KEGG analysis of down- and up-regulated DEGs.**

| Description | Up | Down |
| --- | --- | --- |
| Tryptophan metabolism | 1 | 2 |
| Thiamine metabolism | 2 | 0 |
| Pantothenate and CoA biosynthesis | 0 | 2 |
| Nitrogen metabolism | 0 | 2 |
| Biosynthesis of nucleotide sugars | 2 | 0 |
| Amino sugar and nucleotide sugar metabolism | 3 | 0 |

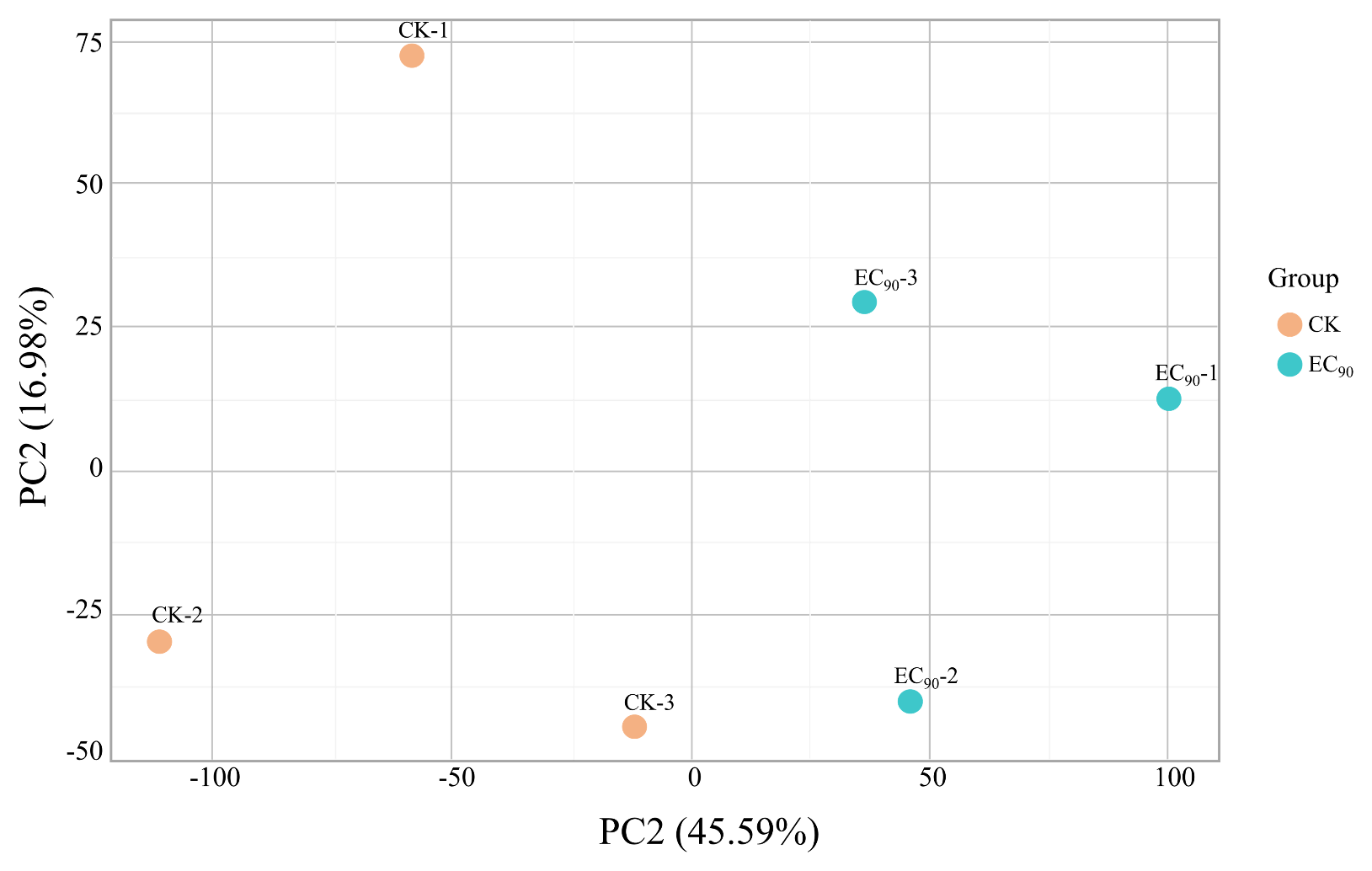

**Fig. S1 Principal component analysis plots between the experimental (EC_90_: 1 μg/mL quinofumelin) and control (CK) samples.**

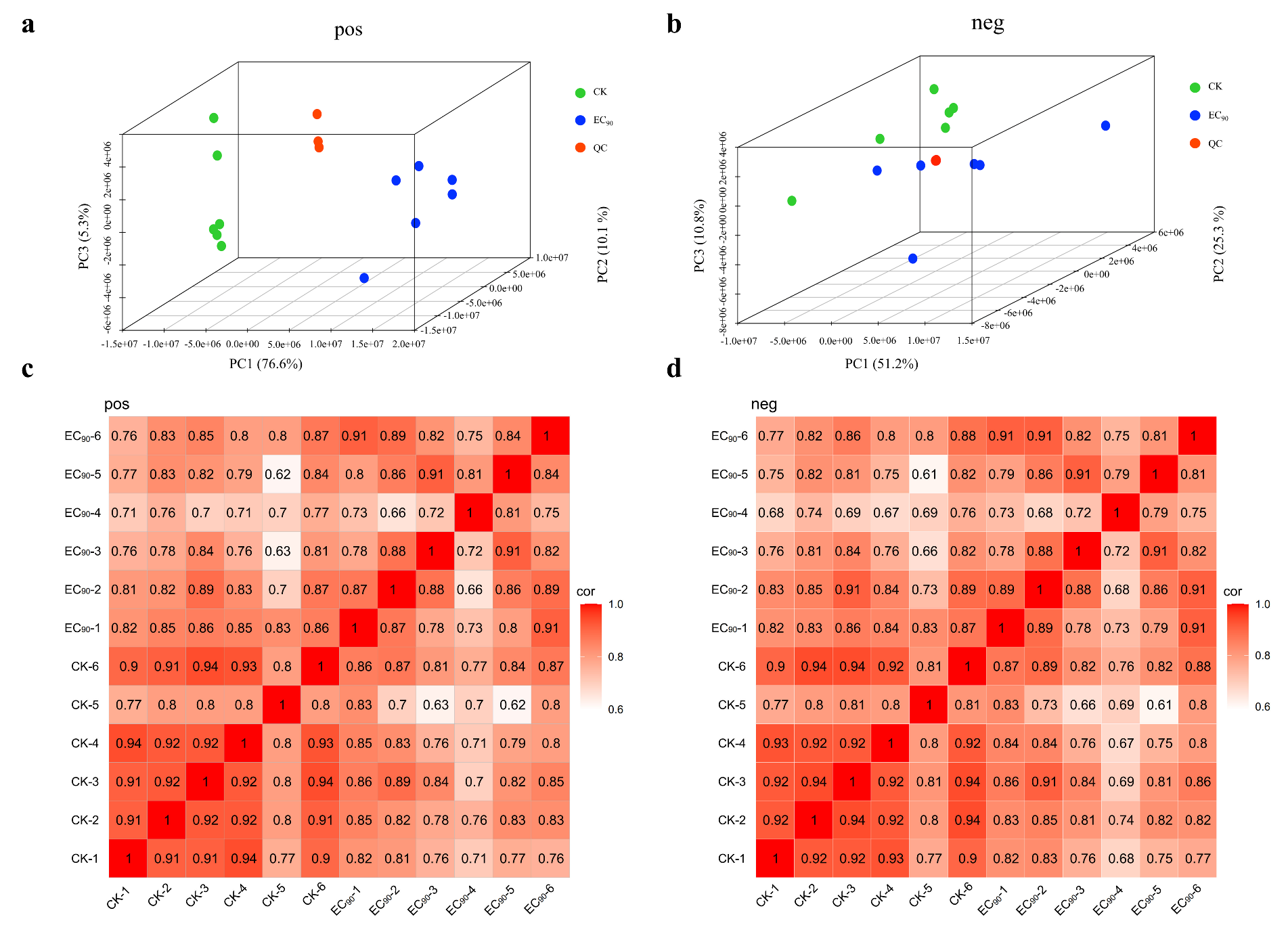

**Fig. S2 Cluster analysis of the metabolite group.**

a 3D principal component analysis diagram of EC_90_, CK and QC samples in positive ion mode.

b 3D principal component analysis diagram of EC_90_, CK and QC samples in negative ion mode.

c Heat map of inter-sample correlation analysis in positive ion mode.

d Heat map of inter-sample correlation analysis in negative ion mode.

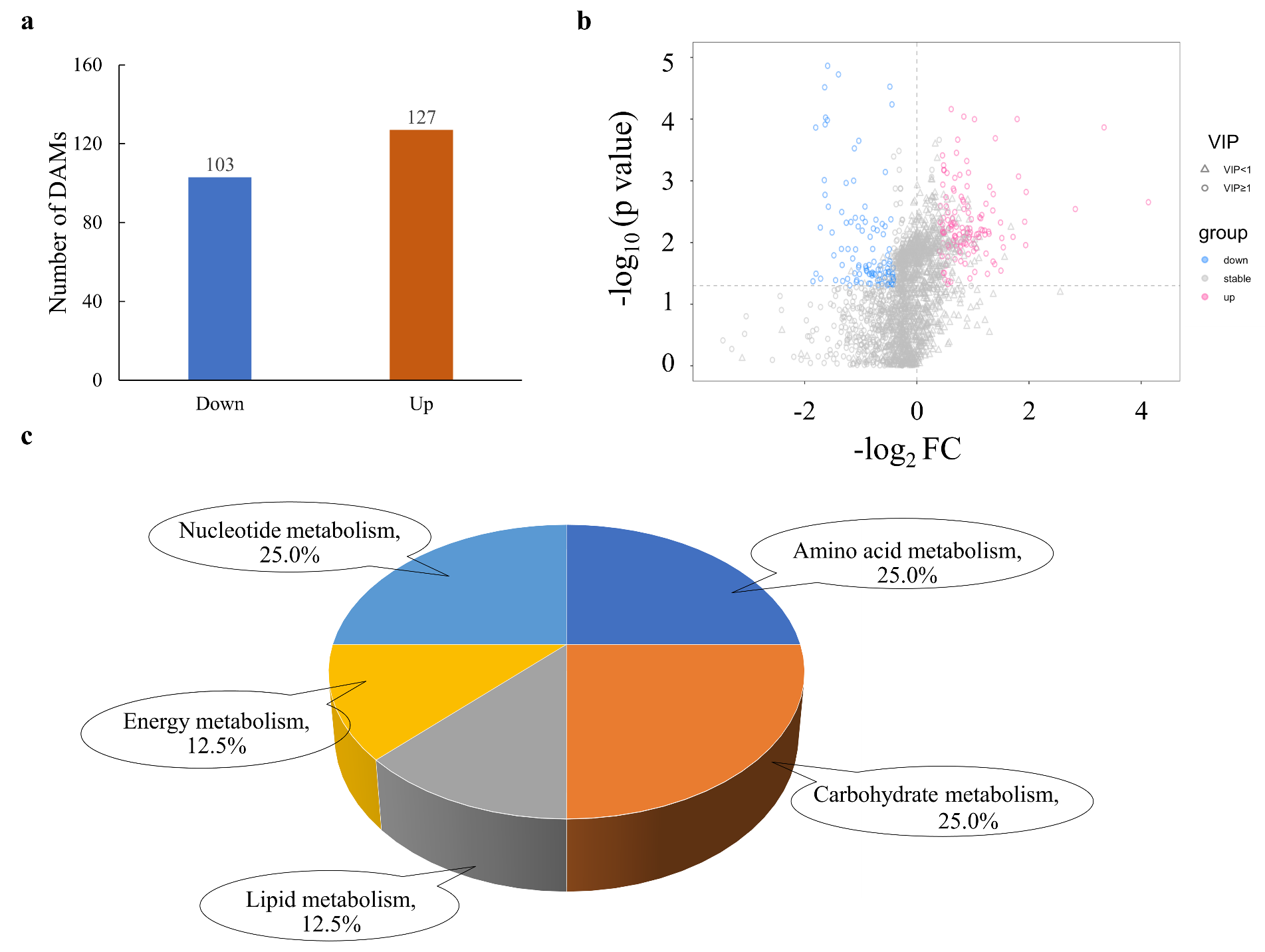

**Fig. S3 Difference analysis of metabolic group data.**

a A Bar chart illustrating the number of DAM quantity in experimental group (EC_90_) and control group (CK) in the negative ion mode.

b Volcanic map of all metabolites in the negative ion mode. The red scatter points represent the up-regulated DAMs, and the blue scatter points represent the down-regulated DAMs.

c Pie diagram of differential metabolite enrichment pathway classification.

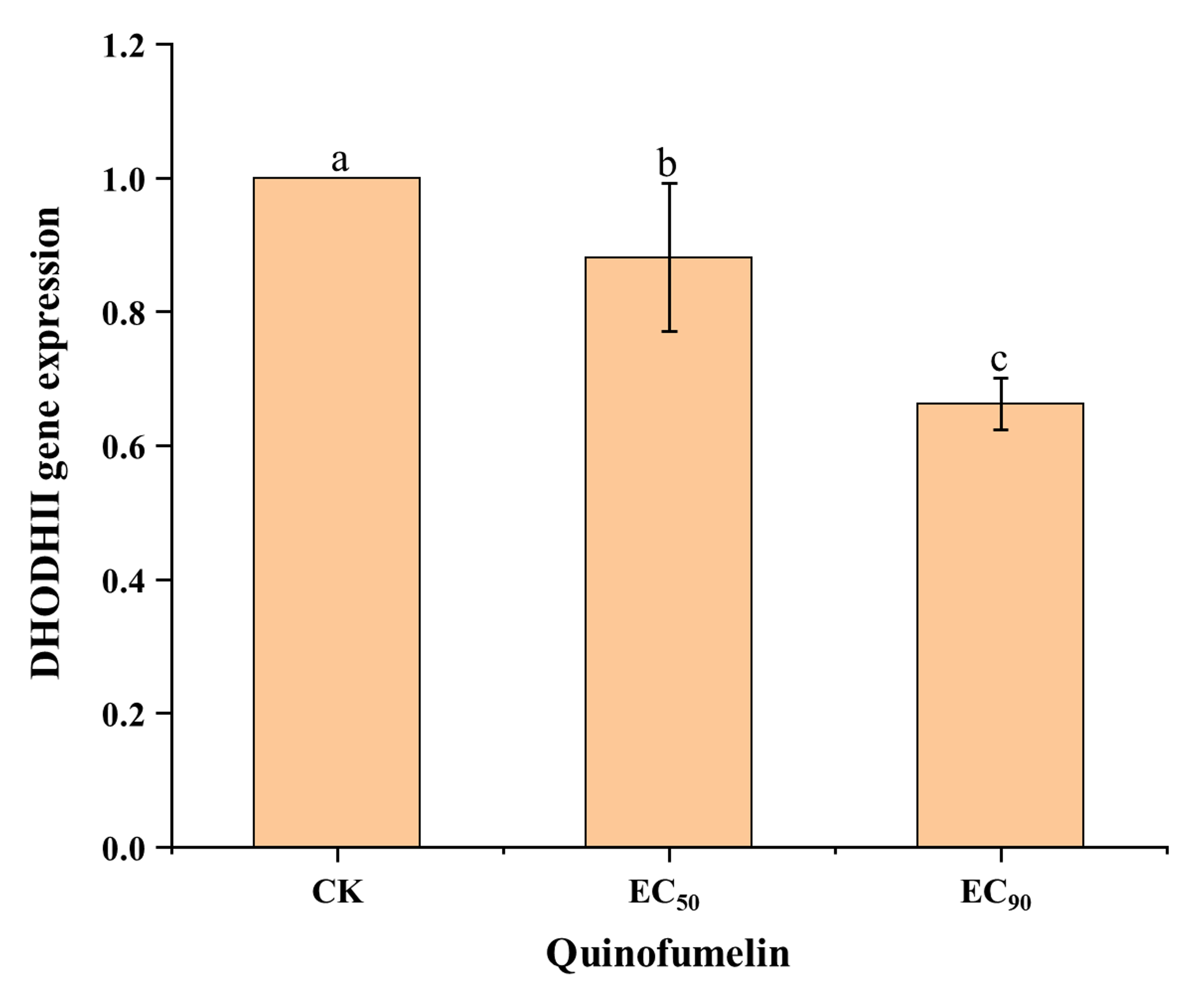

**Fig. S4 DHODHII gene expression in *F. graminearum* affected by quinofumelin.**

After the strain PH-1 was cultured in YEPD medium for 36 h, the EC_50_ (0.035 μg/mL) and EC_90_ (1 μg/mL) of quinofumelin were added respectively, followed by an additional 12-h incubation period. The expression level of the DHODHII gene was subsequently quantified using RT-qPCR. Data represent the mean values and standard errors derived from three independent replicates. Letters above the columns indicate statistically significant differences among treatments (*P*＜0.05, ANOVA, LSD).

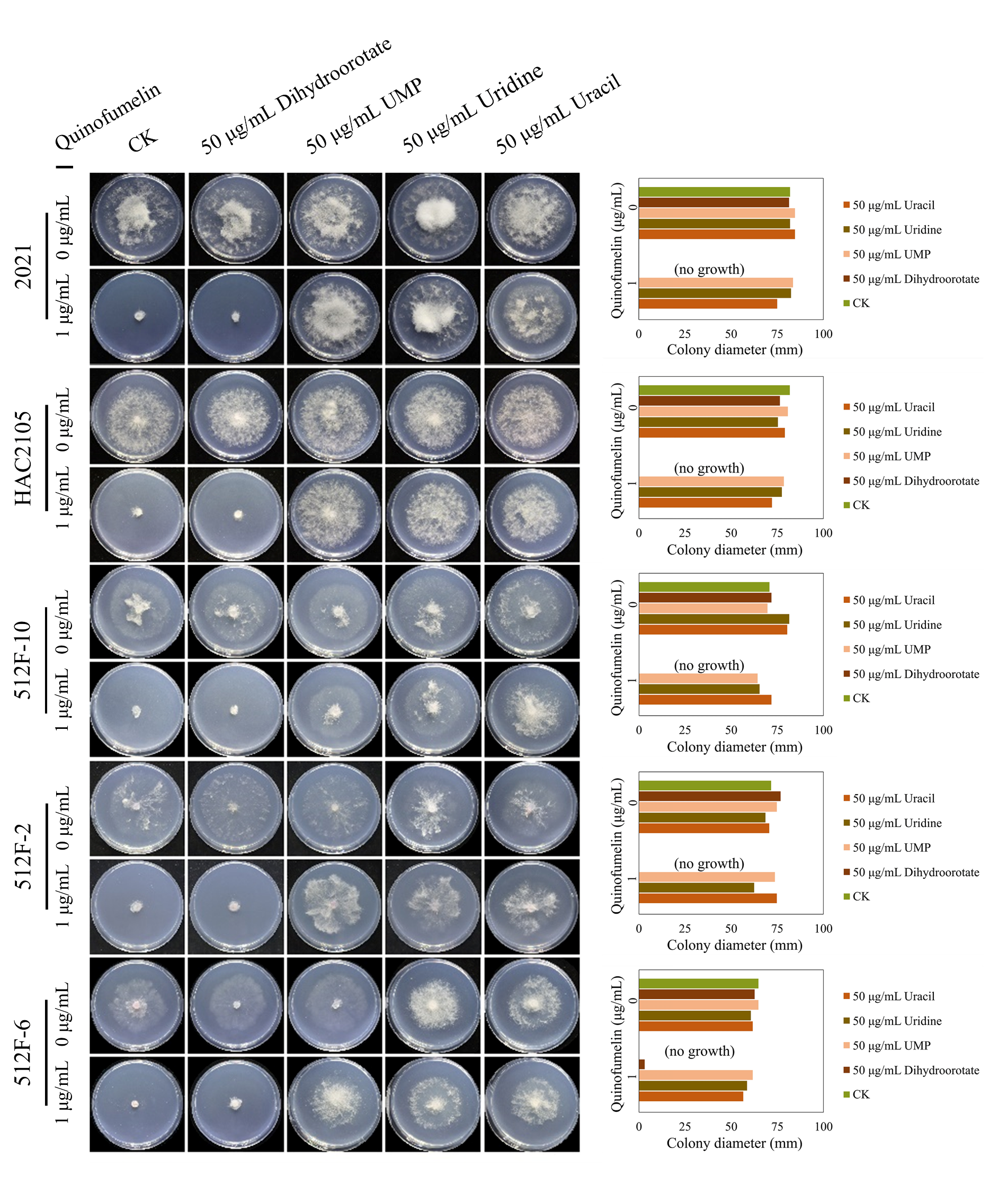

**Fig. S5 Recovery test of mycelial growth suppressed by quinofumelin.**

All strains were incubated on CZA plates at 25°C for 3 days. *F. asiaticum* strains: 2021, HAC2105, 512F-10, 512F-2, and 512F-6. The left image shows the colony morphology, while the right image is a bar chart of colony diameters.

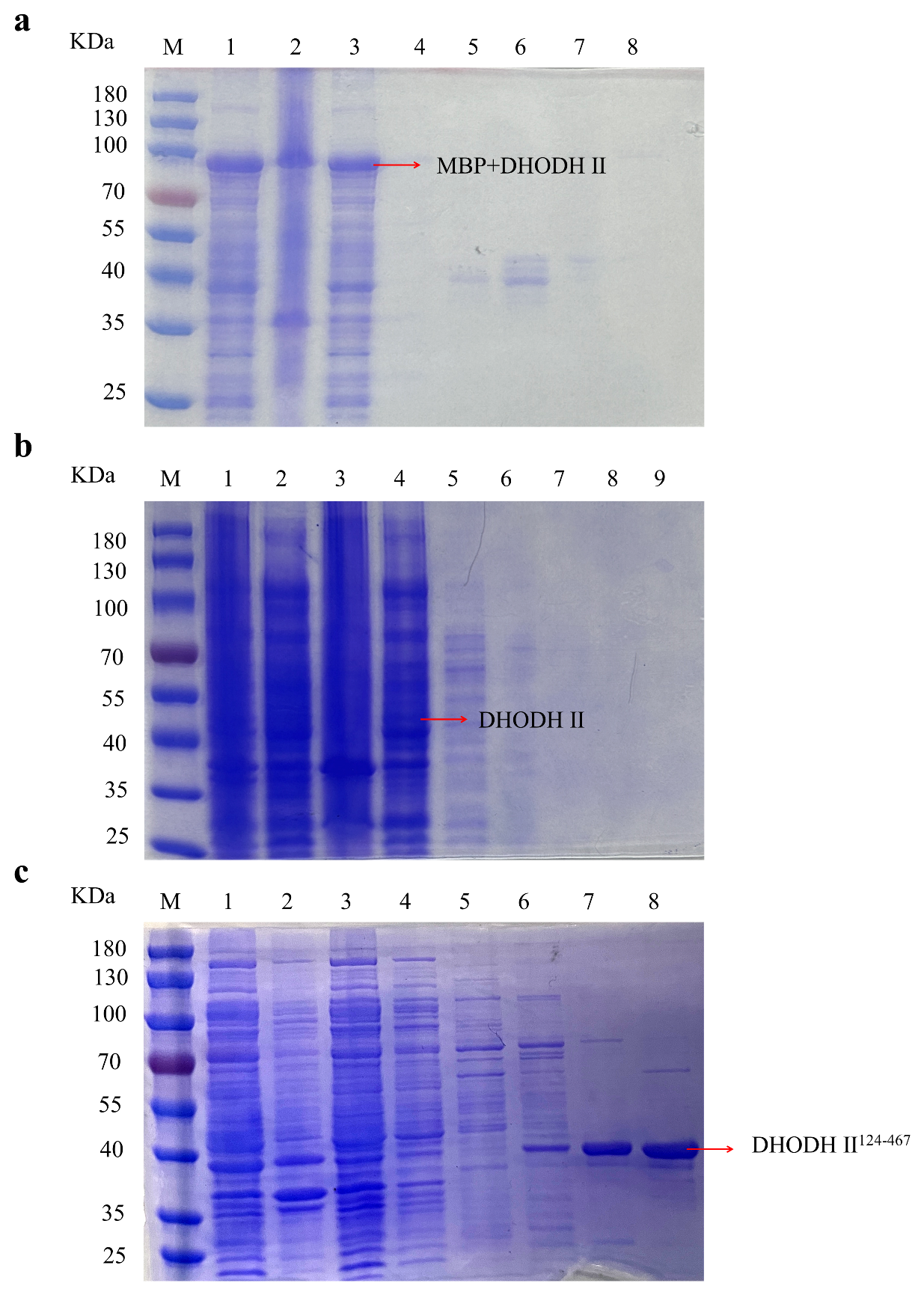

**Fig. S6 SDS-PAGE electrophoresis of the fusion protein.**

a The purification effect of pCold-9×His-MBP-TEV-DHODHII.

b The purification effect of pET-28a (+)-DHODHII.

c The purification of pET-28a (+) -DHODHII^124-467^. The lane M is a protein marker; in a and c, lanes 1-8 were supernatant, precipitate, flow through solution, and eluent containing 5, 20, 50, 100 and 250 mM imidazole, respectively. In b, lanes 1-9 were the whole fungus, supernatant, precipitate, flow-through solution, and eluent containing 5, 20, 50, 100 and 250 mM imidazole.
